# Phylogenomics of free-living neobodonids reveals they are a paraphyletic group from which all other metakinetoplastids are descended

**DOI:** 10.1101/2025.10.03.680260

**Authors:** Daryna Zavadska, Julia A. Packer, Daria Tashyreva, Ryoma Kamikawa, Alastair Simpson, Daniel J. Richter

## Abstract

Kinetoplastea is a major taxon of microbial eukaryotes that includes the well-known trypanosomatid parasites, species of which cause diseases in humans, animals and plants. Free-living kinetoplastids are greatly understudied compared to their parasitic relatives, but are ecologically important microbivores, and collectively comprise the great majority of kinetoplastid diversity. For two decades kinetoplastids have been divided into prokinetoplastids and metakinetoplastids, with the latter further split into four orders, the most diverse of which is Neobodonida. However, the position of the root of the metakinetoplastids and whether neobodonids are a clade has remained unclear due to very limited multi-gene data from free-living kinetoplastids, particularly neobodonids. Here, we present transcriptomic data for eleven newly or recently cultivated free-living neobodonid kinetoplastids. Phylogenomic analyses of a *de novo* generated data set of 444 inferred orthologs and 49 taxa robustly resolve the kinetoplastid tree, including the position of the root of metakinetoplastids. This divides metakinetoplastids such that Trypanosomatida, Eubodonida, Parabodonida, Allobodonidae (formerly neobodonid clade 1E) and neobodonid clade 1D fall on one side, while neobodonid clades Nd6, 1B (Rhynchomonadidae) and 1C fall on the other. Neobodonids are thus inferred to be a paraphyletic group from which all other metakinetoplastids descend. This analysis is the most thorough examination of metakinetoplastid phylogeny thus far, and forms a new basis for tracing the evolutionary history of the entire kinetoplastid group.

**Significance statement:** This study provides transcriptomic data for 11 novel free-living kinetoplastid species, allowing the most accurate phylogeny of metakinetoplastids to date, revealing that neobodonids are paraphyletic and that eubodonids, parabodonids and trypanosomatids emerged within neobodonids. This new framework is essential for understanding the evolution of this major group of parasites and ecologically important free-living heterotrophs.

## Introduction

The kinetoplastids (Kinetoplastea; Euglenozoa) are a major group of heterotrophic protists that are of substantial medical and ecological importance, and that have bizarre basic cell biology: kinetoplast, a mitochondrial genome with massive DNA content, composed largely of thousands of small circular plasmids which encode guide RNAs required for editing of mitochondrial transcripts), paraxonemal rods, intronless nuclear genes transcribed in polycistronic clusters, a lack of transcriptional regulation of nuclear gene expression and glycosomes (modified peroxisomes where glycolysis occurs), among other features. (6; 49; 48; 52; 16; 32). The best studied kinetoplastids are the trypanosomatids, which include agents causing diseases in humans, livestock and plants (51; 20). All other kinetoplastids are historically known as ‘bodonids’. Analyses of sequence data divide kinetoplastids into Prokinetoplastina and Metakinetoplastina (9; 46; 37; 11; 54). Metakinetoplastina includes Trypanosomatida plus three additional major taxa: Eubodonida, Parabodonida and Neobodonida (37; 26). 18S rDNA phylogenies divide neobodonids into subclades ‘1A’, ‘1B’ (Rhynchomonadidae), ‘1C’, ‘Nd6’, ‘1D’, and ‘1E’ (Allobodonidae) (55; 56; 18).

To date, phylogenies 18S rDNA sequences and 1-2 heat shock proteins have not resolved the deep branches within Metakinetoplastina. Unrooted analyses of Metakinetoplastina frequently place eubodonids adjacent to trypanosomatids, generally with poor support (47; 18). Rooted analyses occasionally place some/all neobodonids as the deepest branch within metakinetoplastids (47; 46; 18), indicating that eubodonids, then parabodonids, are successive sisters to trypanosomatids (Figure 1-A). Others place the root elsewhere: within trypanosomatids, or with trypanosomatids sister to all other metakinetoplastids (55; 56; 37) (Figure 1-C). While 18S rDNA datasets have by far the best taxon sampling, an extremely long basal branch separates metakinetoplastids from any outgroup (47; 18; 54). Predictably, the root position is highly unstable and any particular position poorly supported (18). This pattern is less extreme in protein datasets, however taxon sampling is much poorer and the majority of the neobodonid subgroups are missing (46; 13; 58).

**Fig. 1:**
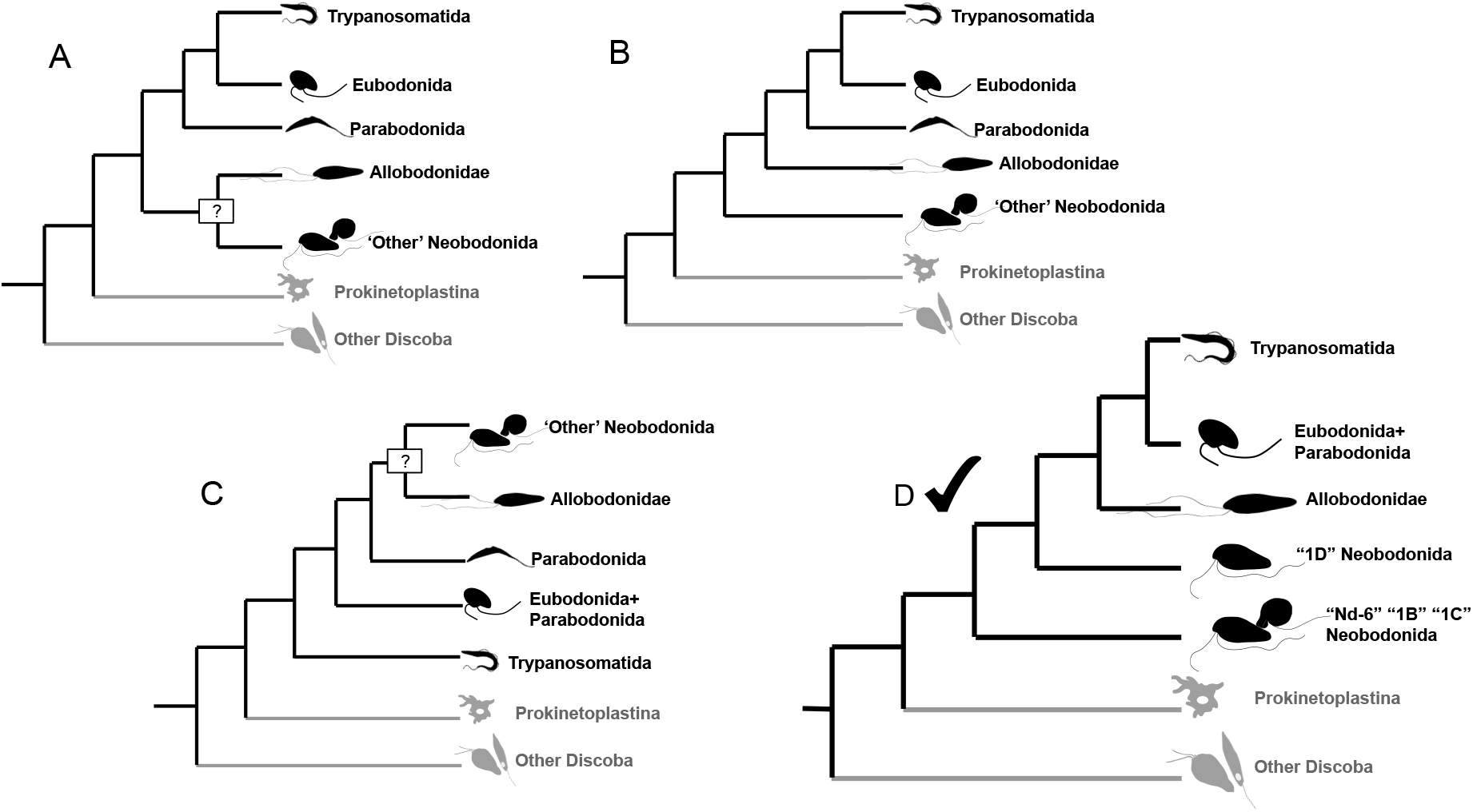
Different hypotheses for the deep phylogeny of kinetoplastids. A-C - various hypotheses on the rooted topology of the phylogeny of the Metakinetoplastina phylogenetic tree outlined in the introduction. D - the topology inferred in this study. “?” indicates an uncertain node in one of the hypothetical topologies.

Resolving the deep-level relationships within Metakinetoplastina is crucial to understand the evolution of the multiple bizarre traits found in kinetoplastids and for a rational high-level systematics of the group.

## Results and Discussion

We sequenced and *de novo* assembled the transcriptomes of 11 neobodonid isolates. *Novijibodo darinka* and *Allobodo yubaba*, were described previously (40), whereas the remaining 9 isolates are undescribed Figure S1, Figure S2. We combined the predicted proteins of these transcriptomes with publicly available datasets representing Metakinetoplastina, Prokinetoplastina and discoban ougroups and performed *de novo* orthogroup inference. We selected single-copy orthogroups (OGs), aligned them individually and concatenated the alignments. The final phylogenomic matrix contained 144,376 sites from 444 *de novo* inferred OGs and 49 species, with an average OG occupancy of 90%, and an average of 86.9% of sites present per species Figure S9.

Maximum Likelihood (ML) and Bayesian analyses recovered identical phylogenies with maximum support at almost all nodes Figure 2; the exceptions were the two internal nodes within Prokinetoplastina, which were poorly resolved in the Bayesian analysis, while the node splitting 1C neobodonids and Rhynchomonadidae had poor support in the ML analysis Figure 2.

**Fig. 2:**
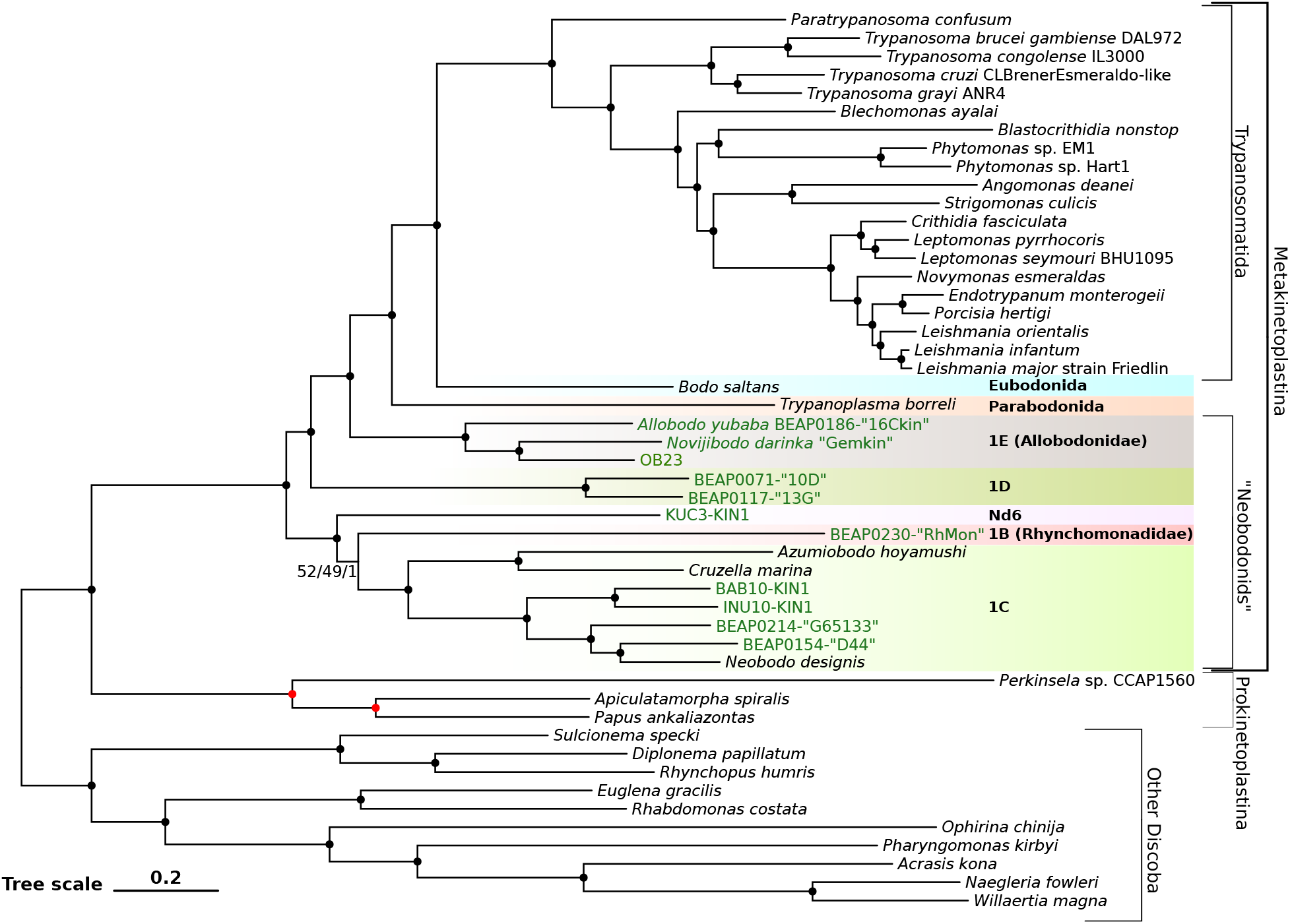
444-protein phylogeny of Kinetoplastea. The topologies of the Maximum Likelihood (ML) and Bayesian phylogenies are identical (apart from internal nodes of Prokinetoplastina, highlighted in red, which are different in the Bayesian phylogeny, see Figure S10). Nodes marked with black circles are recovered in both ML and Bayesian analyses and have 100% support from 1000 ultrafast bootstrap replicates, as well as 200 full bootstrap replicates, and a posterior probability of 1. Nodes without full bootstrap support have ultrafast bootstrap, full bootstrap and posterior probability indicated and separated by slashes. Scale represents expected substitutions per site; branch lengths shown are those of the ML phylogeny inferred under the LG4M+R10 model. Isolates for which the data were produced in this study are highlighted in green. Tree is shown rooted arbitrarily to maximise visual resolution within kinetoplastids.

The monophyly of outgroup lineages (Diplonemida, Euglenida, Heterolobosea) and Trypanosomatida is consistent with previous studies (e.g. (41; 30; 58)), and supports the overall reliability of dataset construction and phylogenomics analysis workflows of our study. Our recovery of Prokinetoplastina and Metakinetoplastina as monophyletic sister taxa confirms previous analyses based on single to several dozens of genes (58; 9; 46; 11; 54).

Previous analyses specifically aiming to resolve deep kinetoplastid phylogeny have included up to two neobodonodis (13; 58). Our taxon sampling covers most of the 18S rDNA-based subgroups of neobodonids for the first time, and arguably represents the first real test of neobodonid monophyly using a phylogenomic approach. Our analyses are in fact consistent with previous studies (13; 58), but the much-improved taxon sampling of neobodonids, and larger number of OGs included (444 versus 64 (13) or 43 (58)) allowed a credible test of neobodonid monophyly, as opposed to inferring their placement under an implicit assumption that they represented a clade. Crucially, we find strong evidence that neobodonids are a paraphyletic assemblage, from which other metakinetoplastids descend as a monophyletic group.

The rooting of Metakinetoplastina splits them into two sister clades that both include ‘neobodonids’. One clade contains Trypanosomatida, Eubodonida, Parabodonida, Allobodonidae (’1E’) and neobodonid clade ’1D’; the second one contains the other three neobodonid clades that were included in the analysis: ‘1C’, ‘Nd6’, and ’1B’ (Rhynchomonadidae). The monophyly of each of ’1E’ (Allobodonidae), ’1D’, and ’1C’ is recovered with full support.

Within the first clade, Eubodonida then Parabodonida are recovered as the closest sister groups to Trypanosomatida, which is consistent with most relevant analyses of hsp90 sequences and with previous phylogenomic analyses (47; 46; 13; 58). The placement of Eubodonida (i.e. *Bodo saltans*) as the closest sister to trypanosomatids has been widely assumed for decades (4; 32), but actually was not clearly resolved by 18S rDNA analyses (47; 37; 18). Our analysis confirms this relationship, however, bolstering previous studies (47; 46; 13; 58).

The placement of Allobodonidae as sister to the Parabodonida, Eubodonida and Trypanosomatida clade is in harmony with the consistent bipartition between Allobodonidae and all other neobodonids in 18S rDNA trees (40; 18; 55; 56). The placement of 1D separate from Nd6, 1C and Rhynchomonadidae in our tree is neither supported nor contraindicated by previous analyses. No prior analyses of protein coding genes included 1D while analyses of 18S rDNA sequences show various sets of poorly supported relationships amongst 1A, Rhynchomonadidae, 1C, 1D and Nd6 (40; 18; 55; 56). The biology of 1D kinetoplastids remains poorly studied. The first isolate from 1D (referred to as ‘*Cryptaulaxoides*-like’) reportedly had a spiral groove running around the cell (55); *Cryptaulaxoides* (= *Cryptaulax* Skuja 1948) refers to elongate, rapidly swimming cells similar to *Rhynchobodo* spp. However, subsequent isolates were listed as ‘*Neobodo* sp.’ (56) and the 1D isolates studied here (BEAP0071-“10D”, BEAP0117-“13G”) superficially resemble *Neobodo designis* (Figure S1, Figure S2).

In the second major clade, the single representative of ’Nd6’ (KUC3-KIN1) falls sister to ’1B’ and ’1C’, although the node grouping ’1B’ and ’1C’ has poor support in the ML analysis. The poor support may be because both Nd6 and 1B are represented by a single species; sampling of more representatives is required. The grouping of Nd6, 1C and 1B is consistent with previous phylogenomic analyses placing the 1B and 1C clades together with strong support to the exclusion of other metakinetoplastids (58). Nd6 placement as a sister to clades 1B and 1C disagrees with some of the formerly produced 18S rDNA phylogenies that showed Nd6 branching with 1C (56) (yet the support in both cases is weak). In other 18S rDNA phylogenies, Nd6 is not recovered as the sister to 1C at all (18).

Our sampling of significant kinetoplastid clades is not exhaustive. The lack of 1A neobodonids is notable, since this clade includes *Rhynchobodo*. Phylogenetic analyses of 1 or 2 heat shock proteins have placed a *Rhynchobodo* isolate separately from other included neobodonids, often as the deepest branch within Metakinetoplastina, albeit statistical support was usually weak, and taxon sampling of both neobodonids and close outgroups was poor (46). If this position is correct, however, it implies that neobodonids are even more serially paraphyletic than our current results already show. A possible deep-branching position for *Rhynchobodo* is intriguing given that the recently documented free-living prokinetoplastids (54), are similar to *Rhynchobodo*: they are polykinetoplastidic cells with tubular extrusomes, large rostrum, a spiral groove or twist to the cell, and tend to swim through fluid rather than along surfaces (5), (39). It might be speculated that the common ancestor of all kinetoplastids was similar to free-living prokinetoplastids and *Rhynchobodo*. However, this might imply multiple independent origins of *N. designis*-like morphotypes: all representatives from 1C, Nd6 and 1D examined in this study are *N. designis*-like Figure S1, Figure S2. Because of this, including neobodonid clade 1A in future analyses would be especially valuable.

Phylogeny, sequences and cultures available from this study will be valuable to address questions on the ancestral states of metakinetoplastid kDNA organisation, glycolysis compartmentalisation, cytoskeleton organisation (as discussed by (47; 32; 2; 40; 59)), metabolic pathways, gene content and gene structure (addressed in (22; 8; 28), but with data from only one to three bodonids), and possible pre-adaptations of ancestral kinetoplastid entities to parasitism.

## Materials and Methods

### Sampling, cultivation and light microscopy

Isolates BEAP0154-“D44”, BEAP0117-“13G” and BEAP0071-“10D” were obtained from plankton samples of subsurface water near Blanes Bay, Mediterranean Sea, Spain (GPS coordinates 41.66667 N, 2.8 E) collected during the year 2022. Isolates BEAP0230-“RhMon” and BEAP0214-“G65133” were obtained from subsurface water of the Caribbean Sea, roughly 10 km south of the island of Grenada (GPS coordinates 11.893732 N, 61.699506 W), in February 2023. We enriched the plankton samples with the nutrient component of RS medium (dilutions from 1:5 to 1:500), and obtained stable mixed protist cultures from the enrichment samples, which were cultivated in RS medium (https://mediadive.dsmz.de/medium/P4, https://mediadive.dsmz.de/medium/P5) with nutrient (NM) to non-nutrient (NNM) component ratios between 1:50 and 1:500, at 17^◦^C (for BEAP0154-“D44”, BEAP0117-“13G” and BEAP0071-“10D”) and at 23^◦^C (BEAP0230-“RhMon” and BEAP0214-“G65133”). The isolates we used in these studies were monoclonal cultures obtained from initial stable mixed cultures by performing 3 to 5 sequential rounds of dilution to extinction. Isolates BEAP0117-“13G”, BEAP0154-“D44”, BEAP0230-“RhMon” and BEAP0214-“G65133” were all initially and routinely cultivated with a 12h:12h light:dark cycle, while isolate BEAP0071-“10D” was cultivated in darkness. We note that, if necessary, all of them are also capable of growth in darkness.

Isolate “OB23” was established from a seawater sample collected at a depth of 5 m at the entrance of Osaka Bay (GPS coordinates 34.324444 N, 135.120833 E), Japan, in July 2023. The sample was enriched in Daigo’s IMK medium (Fujifilm Wako Pure Chemical Corporation). Single cells were then isolated from the enrichment culture using a glass pipette, followed by several sequential rounds of serial dilution to extinction. The established culture was maintained at 20^◦^C under dark conditions in Daigo’s IMK medium.

Freshwater isolates “BAB10-KIN1”, “INU10-KIN1”, and “KUC3-KIN1” were isolated from water samples collected in October 2023 in the Masurian Lakeland, Poland. “BAB10-KIN1” originated from a sample of the hypolimnion of Lake Babiety (53.719288 N, 21.129903 E), while “INU10-KIN1” and “KUC3-KIN1” were isolated from samples of the epilimnion of Lake Inulec (53.805704 N, 21.491791 E) and Lake Kuc (53.815844 N, 21.398909 E), respectively. To enrich for kinetoplastids, 10 mL of each sample were incubated with an autoclaved rice grain for 7 days. Individual kinetoplastid cells were manually picked, together with associated bacteria, using a glass microcapillary and transferred to separate wells of 12-well microplates containing NM component diluted 1:100 in Bold’s Basal Medium with threefold nitrogen and vitamins (3N-BBM (Bischoff, 1963); https://mediadive.dsmz.de/medium/C15). The isolates were routinely maintained in 3N-BBM supplemented with 1:100 dilution of NM at +18^◦^C in the dark.

The isolation of *Novijibodo darinka*-“Gemkin” and *Allobodo yubaba* BEAP0186 “16Ckin” was described previously, in (40).

All isolates in this study were grown as xenic cultures (i.e., with several uncharacterized bacterial species). Light microscopy data from selected isolates used in this study (BEAP0230-“RhMon”, BEAP0214-“G65133”, BEAP0154-“D44”, BEAP0117-“13G”, BEAP0071-“10D”, “BAB10-KIN1”, “INU10-KIN1” and “KUC3-KIN1”) was obtained with a Zeiss Axio Observer.Z1 inverted microscope with a Differential Interference Contrast (DIC) Plan-Apochromat 63x Oil immersion objective (NA=1.4). The imaging setup was equipped with a HDCamC13440-20CU Hamamatsu camera coupled with ZEN image acquisition software. Imaging data was further processed in Fiji (44). Transmission Electron Microscopy was performed at Electron Cryomicroscopy Unit at the Scientific and Technical Centers of the University of Barcelona. Cells were collected by centrifugation (2000 x *g* for 5 min), resuspended in 20% BSA in artificial sea water and cryo-immobilized using a Leica HPM100 high-pressure freezer (Leica Microsystems, Vienna, Austria). Samples were freeze-substituted with 2% osmium tetroxide and 0.1% uranyl acetate dissolved in the pure acetone at -90^◦^C for 72 hours in an EM AFS2 (Leica Microsystems, Vienna, Austria), followed by gradual warming up to RT and infiltration of Epon-812 resin and polymerization in Epon-812 at 60^◦^C for 48 hours. Ultrathin Sections of 60 nm were obtained with a UC6 ultramicrotome (Leica Microsystems, Vienna, Austria) and placed on Formvar-coated copper grids. Sample Sections were stained with 2% uranyl acetate for 30 min, lead citrate for 5 min and examined in a TEM Jeol JEM 1010 (Gatan, Japan) equipped with a tungsten cathode. Images were acquired at 80 kV with a 1k x 1k CCD Megaview III camera. Complete imaging data, including movies, are available via Figshare at the link https://doi.org/10.6084/m9.figshare.30214630.

### RNA isolation and transcriptome sequencing

The *Novijibodo darinka* isolate “Gemkin” was grown to high abundance in a 400 ml parafilm-sealed vented culture flask, initially containing 20 ml of cell culture, 100 ml of CR10 media ((17), with CR number indicating salinity in ppt), and 12 autoclave-sterilised wheat grains. The culture was grown for 12 days in a 21^◦^C dark incubator before being pelleted via centrifugation (1500 x *g* for 12 min, then 6500 x *g* for 3 min). The pelleted cells were lysed using 4 ml TRIzol^TM^ (ThermoFisher), and stored at -20^◦^C for 2 weeks. Total RNA was isolated using the standard TRIzol^TM^ protocol, except that the RNA was precipitated at -20^◦^C overnight. Total RNA was washed twice using the RNeasy PowerClean Pro cleanup kit (QIAGEN).

The RNA was stored at -80^◦^C before being sent to Génome Québec for poly-A selection, cDNA library preparation, and paired-end sequencing (100 bp) using Illumina NovaSeq.

Isolate OB23 was cultivated as described above in eight 50-mL flasks (Violamo) containing Daigo’s IMK medium. After two weeks of cultivation, the cells were pelleted by centrifugation at 2,000 x *g* for 10 min. RNA was extracted from the pelleted cells using TRIzol reagent and Phasemaker Tubes (Thermo Fisher Scientific) according to the manufacturer’s instructions. Total RNA was then used for library construction using the MGIEasy Fast RNA Library Prep Set (MGI Tech). The resulting libraries were circularized using the MGIEasy Dual Barcode Circularization Kit (MGI Tech) and converted into DNA nanoballs (DNBs) using the DNBSEQ-G400RS High-throughput Sequencing Kit and High-throughput Pair-End Sequencing Primer Kit (App-D) (MGI Tech). Finally, 150-bp paired-end sequencing was performed on a DNBSEQ-G400 platform (MGI Tech).

Total RNA from *Allobodo yubaba* (isolate BEAP0186-“16Ckin”), and from the remaining isolates used in this study (BEAP0230-“RhMon”, BEAP0214-“G65133”, BEAP0154-“D44”, BEAP0117-“13G”, BEAP0071-“10D” as well as “BAB10-KIN1”, “INU10-KIN1” and “KUC3-KIN1”) was extracted following the procedure from (42). All of these isolates used in transcriptomics studies were strictly monoclonal (i.e., they were initiated from a single eukaryotic cell and any co-isolated environmental bacteria), except for “INU10-KIN1”, for which the culture originated from a few manually picked cells . For RNA isolation, cultures were grown in RS medium (https://mediadive.dsmz.de/medium/P4, https://mediadive.dsmz.de/medium/P5), of nutrient (NM) to non-nutrient (NNM) component ratios between 1:20 and 1:500 for marine isolates (BEAP0230-“RhMon”, BEAP0214-“G65133”, BEAP0154-“D44”, BEAP0117-“13G”, BEAP0071-“10D”) or 1:100 dilutions of NM component prepared in distilled water in 3N-BBM (Bold’s Basal Medium) (Bischoff, 1963) for freshwater isolates (“BAB10-KIN1”, “INU10-KIN1” and “KUC3-KIN1”), in 75 cm^2^ rectangular canted-neck cell culture flasks with vented caps (353136, Corning Life Sciences) and pelleted by centrifugation (13000 x *g* for 20 min, at +4^◦^C). RNA isolation was performed with an RNAqueous kit (AM1912, ThermoFisher Scientific), followed by genomic DNA digestion using a TURBO DNA-free kit (AM1907, ThermoFisher Scientific). The kits were used according to the manufacturer’s protocols with the modifications described in (42). Double the normal rounds of polyA selection were applied to the total RNA before cDNA synthesis (to reduce the amount of prokaryotic RNA in the sample, which is common in non-axenic cultures). The integrity of total RNA was checked using Bioanalyzer 2100 RNA Pico chips (Agilent Technologies) at the CRG Genomics Core Facility in Barcelona, and total RNA was stored at -80^◦^C before being sent to the same facility for poly-A selection, library preparation, and sequencing. Stranded libraries were prepared using the NEBNext Ultra II Directional RNA Library Prep Kit. Finally, the RNA was sent for paired-end sequencing (150 bp) on an Illumina NextSeq 2000.

### Transcriptome assembly, decontamination, protein prediction and redundancy reduction

The procedure described below and in Figure S3 was performed for all the transcriptomes obtained in this study (isolates “Gemkin”, “OB23”, BEAP0186-“16Ckin”, BEAP0230-“RhMon”, BEAP0214-“G65133”, BEAP0154-“D44”, BEAP0117-“13G”, BEAP0071-“10D”, “BAB10-KIN1”, “INU10-KIN1” and “KUC3-KIN1”).

#### Read trimming and transcriptome assembly

Raw reads were trimmed using fastp(v0.20.1) (12); depending on the sequencing technology used, settings varied (see exact command list at https://github.com/beaplab/kineto_phylo/blob/main/MASTER_commands_V2.sh). Assembly was performed using RNAspades v3.13.1 (7), in stranded mode (with the exception of “OB23”, which was assembled in non-stranded mode), with the other settings left as their defaults.

#### Identification of putative contaminants and creating the database of contaminant genomes

Potential contaminants were primarily identified by searching for 18S ribosomal RNA gene sequences of the non-target eukaryote species and for prokaryotic 16S ribosomal sequences in the assemblies.

18S and 16S of contaminants were retrieved by blasting 18S and 16S sequences of *Mus musculus* (NR 003278) and *Escherichia coli* (NR 114042) against the assembled contigs. Matches were manually filtered out, and target contigs producing reasonable match length, E-value and % identity were selected (see table on FigShare at https://doi.org/10.6084/m9.figshare.30214630 for the exact list of contigs and contaminants). These contigs that putatively contained 18S or 16S of contaminants were used as queries to perform blastn against the NCBI nt database (a custom Perl script was used as a wrapper for performing blastn of multiple queries vs. multiple targets, available on Github https://github.com/beaplab/kineto_phylo/blob/main/Suppl_v1_scripts/web_blast.pl). A single top hit was selected for each contig. NCBI taxonomy IDs of the targets from the top hits were retrieved.

In addition to the taxonomy IDs matching 16S- and 18S-containing contigs identified by the BLAST approach, we also retrieved taxonomy IDs of 18S sequences assembled by phyloFlash v3.3b3 (19) from the raw transcriptomic reads (“Full-length SSU rRNA sequences assembled by SPAdes, matched to SILVA database with Vsearch” list of PhyloFlash output).

Genomes corresponding to all the taxonomy IDs of potential contaminants were downloaded using the Entrez v14.6 and NCBI datasets v16.22.1 tools. If there were multiple genomes available for a single taxonomy ID, up to 5 genomes were selected for download, and RefSeq genomes were given preference.

#### Removal of contamination

For decontamination, assembled contigs were used as blastn queries against the database of putative contaminant genomes obtained in the previous step (a custom Perl script was used as a wrapper for performing blastn of multiple queries vs. multiple targets; it is available on Github https://github.com/beaplab/kineto_phylo/blob/main/Suppl_v1_scripts/run_BLAST.pl).

The distribution of % identity, E-value and match length was visualised for blast hits of 60,000 randomly selected contigs from all assemblies vs. putative contaminant genomes Figure S5 using a custom R script available on Github https://github.com/beaplab/kineto_phylo/blob/main/Suppl_v1_scripts/blast_res_viz.Rmd.

The minimum threshold for blastn hit match length was set to 150 for all assembly-contaminant genome pairs, based on visual inspection of the aforementioned distribution. Minimum thresholds of % identity were selected individually for each pair of assembly and contaminant genome, based on an automated algorithm designed to select a value separating identical or nearly identical matches resulting from contamination versus true matches resulting from sequence homology (see the same custom R script for details).

Contigs that matched contaminant genomes with % identity higher than the automatically identified cutoff and with the length of match exceeding 150 bp, were removed from the corresponding assemblies (using seqkit v0.10.0 (45)).

#### Removal of Cross-contamination

To remove potential cross-contamination among transcriptomes which were sequenced together on the same flow cell (BEAP0230-“RhMon”+BEAP0214-“G65133”, BEAP0186-“16Ckin”+BEAP0154-“D44”+BEAP0117-“13G”+BEAP0071-“10D” and “BAB10-KIN1”+“INU10-KIN1”+“KUC3-KIN1” were sequenced together), the WinstonCleaner https://github.com/kolecko007/WinstonCleaner tool was run with default parameter values. WinstonCleaner removes cross-contamination by mapping trimmed reads from the species sequenced together against the assembled contigs for each species in order to estimate coverage, and when a pair of contigs from different assemblies is found to have a high sequence similarity, the contig with lower read coverage is considered a contaminant and is removed from the corresponding assembly. The cutoffs of minimum % identity for the overlapping contigs to be considered cross-contaminants were automatically identified for each assembly pair and additionally verified by manually inspecting the distribution of % identity of all contigs in assembly pairs.

#### Protein prediction and assembly completeness

Protein prediction was done with the TransDecoder version v5.0.1 tool run in stranded mode (with the exception of “Gemkin” and “OB23”, for which it was run in non-stranded mode) https://github.com/TransDecoder/TransDecoder to produce predicted CDSs and proteins for each decontaminated assembly.

### Obtaining and processing data from other species

Data for other species used in the phylogenomic inference was retrieved from various sources (see table on FigShare at https://doi.org/10.6084/m9.figshare.30214630). Raw reads for *Cruzella marina* (ERR13669942) and *Azumiobodo hoyamushi* (SRR10586159) were downloaded from NCBI and assembled with RNAspades, in stranded mode, with other settings default. Assembled contigs of *Neobodo designis* and *Trypanoplasma/Cryptobia borreli* were obtained from EukProt (43). Assemblies of these four kinetoplastid species (*Neobodo designis, Trypanoplasma/Cryptobia borreli, Cruzella marina, Azumiobodo hoyamushi*) were cleaned using the procedure to remove contamination described above. Assembled contigs of the outgroup species *Pharyngomonas kirbyi* were also obtained from EukProt (43).

For the data from *Tsukubamonas globosa* and *Ploeotia vitrea*, the same procedures were applied as for *Pharyngomonas kirbyi*. However, these two species were excluded from the final set of taxa used in phylogenomic inference, because of low assembly completeness.

Proteins were predicted as described above for *Cruzella marina, Azumiobodo hoyamushi, Neobodo designis, Trypanoplasma/Cryptobia borreli* and *Pharyngomonas kirbyi* (further details are provided in Figure).

Proteomes/CDS from the rest of the species were downloaded from the sources listed in table on FigShare at https://doi.org/10.6084/m9.figshare.30214630, CDSs were translated to amino acid sequences, if necessary, and used in the subsequent procedures in an unaltered state (without additional decontamination or quality control) (see Figure).

### Redundancy reduction

A redundancy-reduced proteome set was generated from the initial set of proteomes using CD-HIT v4.8.1 (31; 15). The redundancy-reduced set (“NR90”) retained proteins with < 90% amino acid global sequence identity for each proteome (the default identity metric is calculated by CD-HIT as the number of aligned identical amino acids divided by the full length of the shorter sequence (31; 15)).

### Orthology inference and verification

The final dataset used for orthology inference contained 11 new predicted proteomes produced in this study, and 38 more predicted proteomes: 28 from kinetoplastid species and 10 from other Discoba lineages (as outgroups), resulting in 49 proteomes in total.

Orthogroups (OGs) were inferred using OrthoFinder v2.5.5 (14), using the “diamond ultra sens” option for the sequence search algorithm and the “msa” option for gene tree inference, with other options left as their defaults.

Initially, 72407 OGs were identified in the set of 49 proteomes. The subset of OGs used for single-OG tree reconstruction and further manual curation was selected based on the species represented and on the number of proteins per species in each OG. Within each OG, we estimated the mean and the standard deviation of the number of proteins present per species, and we counted the number of species with at least one protein. Based on the distributions of these three parameters across all 72407 OGs (see Figure S8), we identified a set of thresholds which would theoretically allow us to produce a subset OGs containing the highest number of single-copy orthologs (i.e., one copy per species) in the largest number of taxa. The summary statistics on the OGs were calculated using a custom R script available on Github https://github.com/beaplab/kineto_phylo/blob/main/Suppl_v1_scripts/Orthofinder_stats_UNIVERSAL_V2.Rmd.

Our thresholds resulted in the selection of OGs meeting all three of the following criteria: (i) mean number of proteins per single species between 0.9 and 1, inclusive, (ii) standard deviation of the number of proteins per single species less than or equal to 0.5, and (iii) number of species with at least one protein present greater than 35 (out of 49 species in total).

The subset obtained by applying these criteria contained 461 OGs. In order to produce a set of single-copy orthologous sequences, first, proteins within each OG were aligned with MAFFT v7.490 (24), using the L-INS-i (“–localpair”) strategy, with the iteration limit set to 16. Resulting alignments were trimmed using trimAl v1.4.rev22 (10) with the “–gappyout” option, and maximum likelihood single-OG trees were inferred using IQ-TREE v1.6.12 (38) with node supports estimated by 1000 ultrafast bootstrap replicates, and other options as their defaults.

Each single-OG tree was manually examined to mark inparalogs, contaminant sequences, and other artefacts for removal from the corresponding OG. As a result of single-gene tree checks, 17 OGs were completely removed from further analysis, 377 OGs were edited by removing some of the proteins, and 67 OGs were left unedited.

Protein sequences from the set of 461 OGs, alignments, single-OG trees, as well as the lists of sequences and OGs that were removed by manual inspection, can be found at table S3 and “orthogroup” directory on FigShare at https://doi.org/10.6084/m9.figshare.30214630.

### Ortholog alignment and trimming

After paralog and artefact removal, proteins of each OG were re-aligned with MAFFT v7.490 (24), using the L-INS-i (“–localpair”) strategy, with the iteration limit set to 1000. Resulting alignments were trimmed using trimAl v1.4.rev22 (10) with the “–gappyout” option. Trimmed alignments of all OGs were concatenated using PhyKit v2.0.1 (50).

### Phylogenomic tree inference

Bayesian tree inference was performed with PhyloBayes-mpi v1.9 (29). Five independent chains were run with the CAT+GTR+G4 model, sampling once every generation. The run was stopped at the 16000th generation, at which point the five chains did not converge, however the only differences between them were in the inter-relationships among the three prokinetoplastids. A consensus tree topology was obtained with a burn-in of 4000 generations (25% of the total number of generations per chain). The mean discrepancy across all bipartitions of all three chains was 0.00631579.

A maximum likelihood analysis was run using IQ-TREE v2.3.6 (36). The LG4M+R10 model was determined by the IQ-TREE ModelFinder (23) as the best-fitting, according to BIC, as well as AIC and AICc. The robustness of the most likely ML tree with the LG4M+R10 model was inferred using ultrafast bootstrapping (1000 replicates) (21) along with standard bootstrapping (200 replicates).

Trees were visualised in tvBOT (57) and edited for presentation with Inkscape.

## Data availability statement

Raw transcriptome sequencing reads for BEAP0186-“16Ckin”, BEAP0230-“RhMon”, BEAP0214-“G65133”, BEAP0154-“D44”, BEAP0117-“13G”, BEAP0071-“10D”, “BAB10-KIN1”, “INU10-KIN1” and “KUC3-KIN1” and “Gemkin” are available via GenBank under the BioProject PRJNA1332963. Raw transcriptome sequencing reads for “OB23” are available via GenBank under the BioProject PRJDB40754. In addition, 18S ribosomal RNA gene sequences obtained from transcriptomic assemblies of BEAP0230-“RhMon”, BEAP0214-“G65133”, BEAP0154-“D44”, BEAP0117-“13G”, and BEAP0071-“10D”, “BAB10-KIN1”, “INU10-KIN1” and “KUC3-KIN1” were deposited in GenBank under IDs PZ229093-PZ229100. The 18S ribosomal RNA gene sequence from “OB23” was deposited in GenBank under IDs LC924540.

A BEAP0186-“16Ckin” cell culture (with mixed bacteria) is available in the Roscoff Culture Collection (RCC) with accession number RCC11336. The following cell cultures (with mixed bacteria) are available in the Culture Collection of Algae and Protozoa (CCAP): BEAP0230-“RhMon” (accession number CCAP 1982/5), BEAP0214-“G65133” (CCAP 1982/4), BEAP0154-“D44” (CCAP 1982/3), BEAP0117-“13G” (CCAP 1982/2), and BEAP0071-“10D” (CCAP 1982/1). All commands that were run during data analysis are stored in the file available on GitHub https://github.com/beaplab/kineto_phylo/blob/main/MASTER_commands_V2.sh. Assembled transcriptome contigs and predicted proteins for all newly sequenced isolates in this study are available on FigShare at https://doi.org/10.6084/m9.figshare.30214630.

All files generated in the process that cannot be directly obtained by running the commands from the aforementioned files (for example, the manually curated sets of orthologous genes) are available either through the GitHub repository https://github.com/beaplab/kineto_phylo or Figshare at https://doi.org/10.6084/m9.figshare.30214630.

## Acknowledgments

This project has received funding from the European Research Council (ERC) under the European Union’s Horizon 2020 research and innovation programme (grant agreement No. 949745), from the Departament de Recerca i Universitats de la Generalitat de Catalunya (exp. 2021 SGR 00751), from grant PID2023-152955NA-I00 funded by MICIU/AEI/10.13039/501100011033 and by ERDF/EU (to D.J.R.), from the Natural Sciences and Engineering Research Council of Canada (NSERC) discovery grant 298366-2019 (to A.G.B.S.), from Grants-in-Aid for Scientific Research (No. 21H05057, No. 24K21929, and No. 25K02334) from the Japan Society for the Promotion of Science (JSPS), from the project No. 2022/47/P/NZ8/02074 (to DT) cofunded by the National Science Centre and the European Union’s Horizon 2020 research and innovation programme under the Marie Sklodowska-Curie grant agreement no. 945339. RK is also supported by Institute for Fermentation, Osaka (2024-G-1-011); some of the environmental samples were collected thanks to funding from the National Science Centre (Poland), grant number 2020/37/B/NZ8/01456 (to Anna Karnkowska).

Computational analyses were enabled by the supercomputing infrastructure from the Galician Supercomputing Center (CESGA; funded by the Spanish Ministry of Science and Innovation, the Galician Government and the European Regional Development Fund; ERDF).

We are deeply grateful to Clare Morrall for arranging access to sampling and facilities in Grenada. We thank Clara Cardelús, operating the Blanes Bay Microbial Observatory (BBMO), and Josep M. Gasol and Ramon Massana for providing access to monthly water samples from Blanes Bay. We thank Margarita Skamnelou and À lex Gàlvez i Morante from the Biology and Ecology of Abundant Protists lab for isolating, identifying and establishing initial cultures of isolates of BEAP0154-“D44”, BEAP0230-“RhMon” and BEAP0214-“G65133”. We thank Dr. Keigo Yamamoto (Research Institute of Environment, Agriculture and Fisheries, Osaka Prefecture, Osaka, Japan) for arranging access to sampling and facilities in Osaka Bay. We thank Anna Karnkowska for providing environmental samples from which isolates “BAB10-KIN1”, “INU10-KIN1”, and “KUC3-KIN1” were isolated. We also thank Elizabeth Weston (Dalhousie University) for training JP in RNA extraction and maintaining cultures.

## References

Adl SM et al. 2019. Revisions to the classification, nomenclature, and diversity of eukaryotes. Journal of Eukaryotic Microbiology. 66:4–119. doi: 10.1111/jeu.12691

Andrade-Alviárez D et al. 2022. Delineating transitions during the evolution of specialised peroxisomes: Glycosome formation in kinetoplastid and diplonemid protists. Frontiers in Cell and Developmental Biology. 10:979269. doi: 10.3389/fcell.2022.979269

Bischoff, H. 1963. Some soil algae from Enchanted Rock and related algal species. Phycological Studies. IV

Blom, D et al. 1998. RNA editing in the free-living bodonid Bodo saltans. Nucleic Acids Research. 26:1205–1213

Brugerolle G. 1985. Des trichocystes chez les bodonides, un caractère phylogénétique supplémentaire entre kinetoplastida et euglenida. Protistologica (Paris). 21:339–348.

Brugerolle G, Lom J, Nohynkova E, Joyon L. 1979. Comparaison Et evolution Des structures cellulaire chez plusieurs especes de Bodonides et Cryptobiides appartenant aux genres Bodo, Cryptobia et Trypanoplasma (Kinetoplastida, Mastigophora). Protistologica. 15:197–221.

Bushmanova E, Antipov D, Lapidus A, Prjibelski AD. 2019. RnaSPAdes: a de novo transcriptome assembler and its application to RNA-Seq data. GigaScience. 8. doi: 10.1093/gigascience/giz100.

Butenko A et al. 2020. Evolution of metabolic capabilities and molecular features of diplonemids, kinetoplastids, and euglenids. BMC Biology. 18. doi: 10.1186/s12915-020-0754-1.

Callahan HA, Litaker RW, Noga EJ. 2002. Molecular Taxonomy of the Suborder Bodonina (Order Kine-toplastida), Including the Important Fish Parasite, Ichthyobodo necator. Journal of Eukaryotic Microbiology. 49:119–128. doi: 10.1111/j.1550-7408.2002.tb00354.x.

Capella-Gutierrez S, Silla-Martinez JM, Gabaldon T. 2009. TrimAl: a tool for automated alignment trimming in large-scale phylogenetic analyses. Bioinformatics. 25:1972–1973. doi: 10.1093/bioinformatics/btp348.

Cenci U et al. 2016. Heme pathway evolution in kinetoplastid protists. BMC Evolutionary Biology. 16. doi: 10.1186/s12862-016-0664-6.

Chen S, Zhou Y, Chen Y, Gu J. 2018. Fastp: an ultra-fast all-in-one FASTQ preprocessor. Bioinformatics. 34:i884–i890. doi: 10.1093/bioinformatics/bty560.

Deschamps P et al. 2010. Phylogenomic analysis of kinetoplastids supports that trypanosomatids arose from within bodonids. Molecular Biology and Evolution. 28:53–58. doi: 10.1093/molbev/msq289.

Emms DM, Kelly S. 2019. OrthoFinder: phylogenetic orthology inference for comparative genomics. Genome Biology. 20. doi: 10.1186/s13059-019-1832-y.

Fu L, Niu B, Zhu Z, Wu S, Li W. 2012. CD-HIT: accelerated for clustering the nextgeneration sequencing data. Bioinformatics. 28:3150–3152. doi: 10.1093/bioinformatics/bts565.

Gibson W. 2017. Handbook of the Protists. Springer International Publishing. doi: 10.1007/978-3-319-32669-6.

Gigeroff AS, Eglit Y, Simpson AGB. 2023. Characterisation and Cultivation of New Lineages of Colponemids, a Critical Assemblage for Inferring Alveolate Evolution. Protist. 174:125949. doi: 10.1016/j.protis.2023.125949.

Goodwin JD, Lee TF, Kugrens P, Simpson AGB. 2018. Allobodo chlorophagus n. gen. n. sp., a Kinetoplastid that Infiltrates and Feeds on the Invasive Alga Codium fragile. Protist. 169:911–925. doi: 10.1016/j.protis.2018.07.001.

Gruber-Vodicka HR, Seah KB, Pruesse E. 2020. phyloFlash: Rapid Small-Subunit rRNA Profiling and Targeted Assembly from Metagenomes. MSystems. 5. doi: 10.1128/msystems.00920-20.

Hide G. 1999. History of Sleeping Sickness in East Africa. Clinical Microbiology Reviews. 12:112–125. doi: 10.1128/cmr.12.1.112.

Hoang DT, Chernomor O, von Haeseler A, Minh BQ, Vinh LS. 2017. UFBoot2: Improving the Ultrafast Bootstrap Approximation. Molecular Biology and Evolution. 35:518–522. doi: 10.1093/molbev/msx281.

Jackson, Andrew P and Otto Thomas D and Aslett, Martin and Armstrong Stuart D and Bringaud, Frederic and Schlacht, Alexander and Hartley, Catherine and Sanders, Mandy and Wastling Jonathan M and Dacks Joel B and others. 2016. Kinetoplastid phylogenomics reveals the evolutionary innovations associated with the origins of parasitism. Current Biology 26.2:161–172. doi: 10.1016/j.cub.2015.11.055.

Kalyaanamoorthy S, Minh BQ, Wong TKF, von Haeseler A, Jermiin LS. 2017. ModelFinder: fast model selection for accurate phylogenetic estimates. Nature Methods. 14:587–589.

Katoh K, Standley DM. 2013. MAFFT Multiple Sequence Alignment Software Version 7: Improvements in Performance and Usability. Molecular Biology and Evolution. 30:772–780. doi: 10.1093/molbev/mst010.

Koch TA, Ekelund F. 2005. Strains of the Heterotrophic Flagellate Bodo designis from Different Environments Vary Considerably with Respect to Salinity Preference and SSU rRNA Gene Composition. Protist. 156:97–112. doi: 10.1016/j.protis.2004.12.001.

Kostygov AY et al. 2021. Euglenozoa: taxonomy, diversity and ecology, symbioses and viruses. Open Biology. 11. doi: 10.1098/rsob.200407.

Kostygov AY et al. 2024. Phylogenetic framework to explore trait evolution in Trypanosomatidae. Trends in Parasitology. 40:96–99. doi: 10.1016/j.pt.2023.11.009.

Kostygov AY et al. 2024. Comprehensive analysis of the Kinetoplastea intron landscape reveals a novel intron-containing gene and the first exclusively trans-splicing eukaryote. BMC biology. 22:281. doi: 10.1186/s12915-024-02080-z

Lartillot N, Rodrigue N, Stubbs D, Richer J. 2013. PhyloBayes MPI: Phylogenetic Reconstruction with Infinite Mixtures of Profiles in a Parallel Environment. Systematic Biology. 62:611–615. doi: 10.1093/sysbio/syt022.

Lax G et al. 2021. Multigene phylogenetics of euglenids based on single-cell transcriptomics of diverse phagotrophs. Molecular Phylogenetics and Evolution. 159:107088. doi: 10.1016/j.ympev.2021.107088.

Li W, Godzik A. 2006. CD-hit: a fast program for clustering and comparing large sets of protein or nucleotide sequences. Bioinformatics. 22:1658–1659. doi: 10.1093/bioinformatics/btl158.

Lukeš J et al. 2002. Kinetoplast DNA Network: Evolution of an Improbable Structure. Eukaryotic Cell. 1:495–502. doi: 10.1128/ec.1.4.495-502.2002.

Lukeš J et al. 2018. Trypanosomatids Are Much More than Just Trypanosomes: Clues from the Expanded Family Tree. Trends in Parasitology. 34:466–480. doi: 10.1016/j.pt.2018.03.002.

Lukeš J et al. 2023. Trypanosomes as a magnifying glass for cell and molecular biology. Trends in Parasitology. 39:902–912. doi: 10.1016/j.pt.2023.08.004.

Manni M, Berkeley MR, Seppey M, Simão FA, Zdobnov EM. 2021. BUSCO Update: Novel and Streamlined Workflows along with Broader and Deeper Phylogenetic Coverage for Scoring of Eukaryotic, Prokaryotic, and Viral Genomes Kelley, J, editor. Molecular Biology and Evolution. 38:4647–4654. doi: 10.1093/molbev/msab199.

Minh BQ et al. 2020. IQ-TREE 2: New Models and Efficient Methods for Phylogenetic Inference in the Genomic Era. Molecular Biology and Evolution. 37:1530–1534. doi: 10.1093/molbev/msaa015.

Moreira D, López-García P, Vickerman K. 2004. An updated view of kinetoplastid phylogeny using environmental sequences and a closer outgroup: proposal for a new classification of the class Kinetoplastea. International Journal of Systematic and Evolutionary Microbiology. 54:1861–1875. doi: 10.1099/ijs.0.63081-0.

Nguyen L-T, Schmidt HA, von Haeseler A, Minh BQ. 2014. IQ-TREE: A Fast and Effective Stochastic Algorithm for Estimating Maximum-Likelihood Phylogenies. Molecular Biology and Evolution. 32:268–274. doi: 10.1093/molbev/msu300.

Nikolaev SI et al. 2003. The taxonomic position of Klosteria bodomorphis gen. and sp. nov. (Kinetoplastida) based on ultrastructure and SSU rRNA gene sequence analysis. Protistology. 3:126–135. doi:

Packer JA et al. 2025. Characterization of Allobodo yubaba sp. nov. and Novijibodo darinka gen. et sp. nov., cultivable free-living species of the phylogenetically enigmatic kinetoplastid taxon Allobodonidae. Journal of Eukaryotic Microbiology. 72. doi: 10.1111/jeu.13072.

Pánek T et al. 2025. An expanded phylogenomic analysis of Heterolobosea reveals the deep relationships, non-canonical genetic codes, and cryptic flagellate stages in the group. Molecular Phylogenetics and Evolution. 204:108289–108289. doi: 10.1016/j.ympev.2025.108289.

Richter DJ, Parinaz Fozouni, Eisen MB, King N. 2018. Gene family innovation, conservation and loss on the animal stem lineage. eLife. 7. doi: 10.7554/elife.34226.

Richter DJ et al. 2022. EukProt: A database of genome-scale predicted proteins across the diversity of eukaryotes. Peer Community Journal. 2. doi: 10.24072/pcjournal.173.

Schindelin J et al. 2012. Fiji: an open-source Platform for biological-image Analysis. Nature Methods. 9:676–82.

Shen W, Le S, Li Y, Hu F. 2016. SeqKit: A Cross-Platform and Ultrafast Toolkit for FASTA/Q File Manipulation. PLOS ONE. 11:e0163962. doi: 10.1371/journal.pone.0163962.

Simpson, AGB et. al. 2004. Early Evolution within Kinetoplastids (Euglenozoa), and the Late Emergence of Trypanosomatids. Protist. 155:407–422. doi: 10.1078/1434461042650389.

Simpson AGB, Lukeš J, Roger AJ. 2002. The Evolutionary History of Kinetoplastids and Their Kinetoplasts. Molecular Biology and Evolution. 19:2071–2083. doi: 10.1093/oxfordjournals.molbev.a004032.

Simpson AGB, Stevens JR, Lukeš J. 2006. The evolution and diversity of kinetoplastid flagellates. Trends in Parasitology. 22:168–174. doi: 10.1016/j.pt.2006.02.006.

Simpson AGB. 1997. The identity and composition of the Euglenozoa. Archiv Fur Protistenkunde. 148:318–328. doi: 10.1016/s0003-9365(97)80012-7.

Steenwyk JL et al. 2021. PhyKIT: a broadly applicable UNIX shell toolkit for processing and analyzing phylogenomic data Schwartz, R, editor. Bioinformatics. 37:2325–2331. doi: 10.1093/bioinformatics/btab096.

Steverding D. 2008. The history of African trypanosomiasis. Parasites & Vectors. 1:3. doi: 10.1186/1756-3305-1-3.

Tashyreva D et al. 2022. Diplonemids – A Review on ‘New’ Flagellates on the Oceanic Block. Protist. 173:125868–125868. doi: 10.1016/j.protis.2022.125868.

Tice AK et al. 2021. PhyloFisher: A phylogenomic package for resolving eukaryotic relationships. PLOS Biology. 19:e3001365–e3001365. doi: 10.1371/journal.pbio.3001365.

Tikhonenkov DV, Ryan M.R. Gawryluk, Mylnikov AP, Keeling PJ. 2021. First finding of free-living representatives of Prokinetoplastina and their nuclear and mitochondrial genomes. Scientific Reports. 11. doi: 10.1038/s41598-021-82369-z.

Von der Heyden S, Chao E, Vickerman K Cavalier-Smith. 2004. Ribosomal RNA Phylogeny of Bodonid and Diplonemid Flagellates and the Evolution of Euglenozoa. The Journal of Eukaryotic Microbiology. 51:402–416. doi: 10.1111/j.1550-7408.2004.tb00387.x.

Von der Heyden S, Cavalier-Smith T. 2005. Culturing and environmental DNA sequencing uncover hidden kinetoplastid biodiversity and a major marine clade within ancestrally freshwater Neobodo designis. International Journal of Systematic and Evolutionary Microbiology. 55:2605–2621. doi: 10.1099/ijs.0.63606-0.

Xie J et al. 2023. Tree Visualization By One Table (tvBOT): a web application for visualizing, modifying and annotating phylogenetic trees. Nucleic Acids Research. 51:W587–W592. doi: 10.1093/nar/gkad359

Yazaki E et al. 2017. Global Kinetoplastea phylogeny inferred from a large-scale multigene alignment including parasitic species for better understanding transitions from a free-living to a parasitic lifestyle. Genes & Genetic Systems. 92:35–42. doi: 10.1266/ggs.16-00056.

Yubuki N, Čepička I. and Leander BS. 2016. Evolution of the microtubular cytoskeleton (flagellar apparatus) in parasitic protists. Molecular and biochemical parasitology. 209:26–34. doi: 10.1016/j.molbiopara.2016.02.002.

