## Supplemental material for "Phylogenomics of free-living neobodonids reveals they are a paraphyletic group from which all other metakinetoplastids are descended" (from another version of this preprint: version 2, 2026-04-09)

### Supplemental information

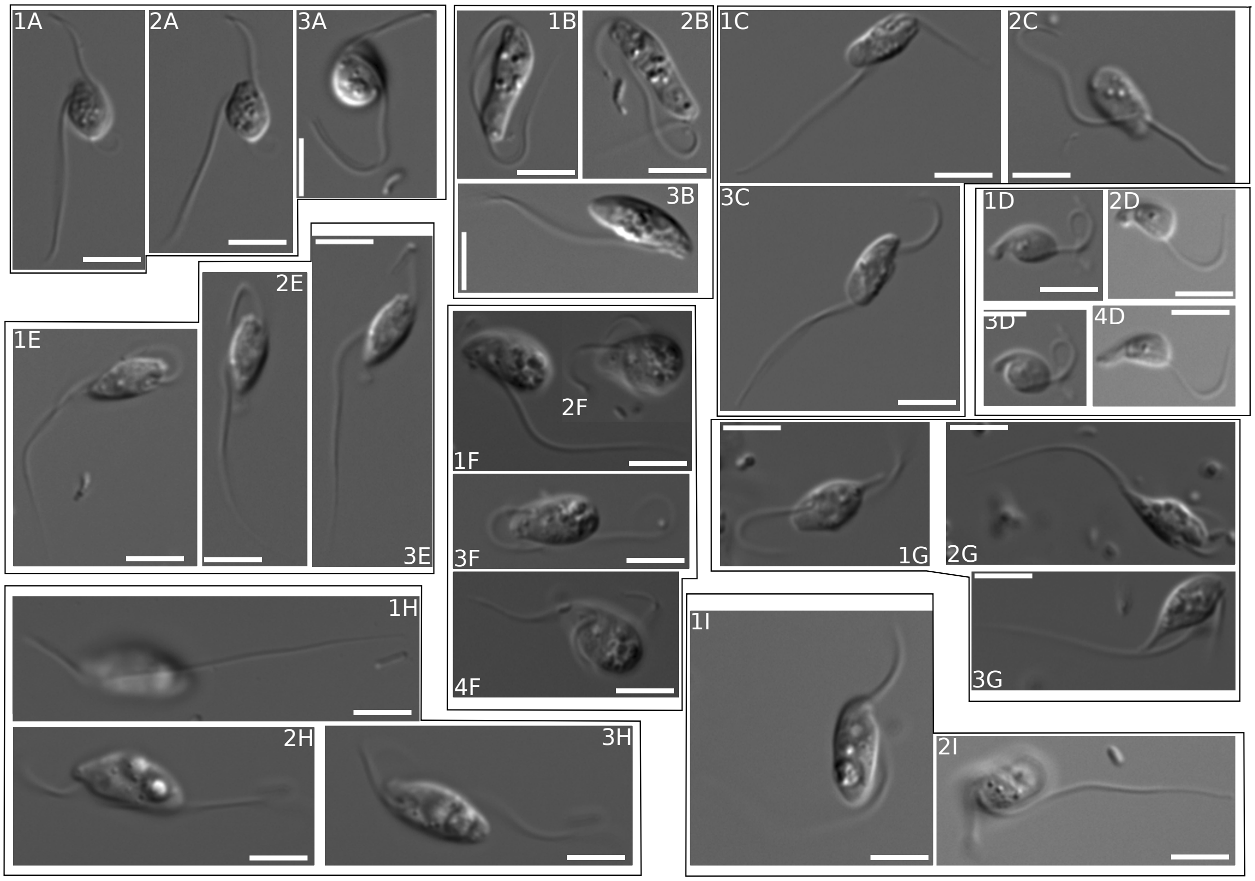


Figure Figure S1: **DIC images of isolates from this study.** The letters correspond to the isolates, and numbers represent different images of the same isolate. A - BEAP0071-"10D", B - BEAP0117-"13G", C - BEAP0154-"D44", D - BEAP0230-"RhMon", E - BEAP0214-"G65133", F - "KUC3-KIN1", G - "OB23", H - "BAB10-KIN1", I - "INU10-KIN1". Scale bar is 5 μm in all images. All live DIC imaging data is available on FigShare `https://doi.org/10.6084/m9.figshare.30214630`

#### S1.1 Selecting settings and dataset for Orthology inference

Various combinations of (i) sets of taxa used in orthology inference, (ii) redundancy reduction algorithms applied to the proteomes prior to the orthology inference, and (iii) settings of OrthoFinder software were tested before selecting the combination used for phylogenomic analysis in the main text. The specific values that we tested in combination were: (i) sets of 40, 45, 48, 50 and 52 taxa (to see which taxa were included, see table on FigShare at `https://doi.org/10.6084/m9.figshare.30214630`). The sets of taxa were formed based on the occurrence of genes from each species in OGs Figure S7, and based on the taxonomic identity of species. Species from Kinetoplastea were kept regardless of their representation across OGs. In order to select suitable outgroup species, an excess of outgroup taxa was included in the largest 52-species dataset (see S2.2 below). As some of these datasets were partial, they were principally those which were removed in the reduced taxa sets (40-50 taxa). (ii) redundancy-reduced proteomes which retained sequences of <90%, or <97% amino acid global sequence identity and (iii) Orthofinder settings: the MCL inflation parameter ("-I" option), the sequence search program ("-S" option) and the method for gene tree inference ("-M" option). The combinations of settings and datasets tested are presented in table S3 on FigShare at `https://doi.org/10.6084/m9.figshare.30214630`. This set of combinations is not exhaustive, but we assume that some settings could be excluded based on the performance of one combination. The OGs inferred from each combination of settings and datasets were filtered by establishing the following set of threshold values: (a) mean number of proteins per single species between 0.9 and 1, inclusive, (b) standard deviation of the number of proteins per single species less than or equal to 0.5, and (c) number of species with at least one protein present greater than 35 (out of 45 species in total). For the resulting subsets of OrthoFinder OGs produced by each combination of settings i-iii, we evaluated performance by assessing the number of OGs fulfilling all three criteria a-c Figure S6, as well as the representation of each species across the OGs in the subset Figure S7.

#### S2.2 Selecting species for phylogenomics dataset

The initial dataset included:

- All species from "bodonids" (non-Trypanosomatid Metakinetoplastina, 5 species) and Prokinetoplastina (3 species) for which data was publicly available, and 11 proteomes of "bodonid" species generated in this study.
- A subset of 20 representatives of Trypanosomatida. Since well-annotated proteomes from multiple closely-related species of Trypanosomatida are available in excess, a subset was selected, taking (i) up to 4 species from a single genus; if more than 4 species per genus were available, and the (ii) phylogenetic position of a species within a genus and (iii) quality of proteome annotation were taken into account.
- Outgroup, which initially contained a set of 17 species from other lineages of Discoba. The diversity of the outgroup species was limited to Discoba (rather than expanded with species from other supergroups) because the OGs were inferred *de novo*, and unnecessarily increasing the diversity of the outgroup might have led to a reduction in the number of single-copy orthologs, since many OGs could be Discoba-specific.

7 species initially selected to be a part of the outgroup were discarded after several initial attempts of *de novo* OG inference, as their genes were missing from the majority of inferred OGs (and Euglenozoa-specific BUSCO completeness scores were also low for these species).

| | *De novo* assembly | Decontamination | Protein prediction | Downstream (redundancy reduction and on) |
|---|---|---|---|---|
| *Cruzella marina* | X | X | X | X |
| *Azumiobodo hoyamushi* | X | X | X | X |
| *Neobodo designis* | | X | X | X |
| *Trypanoplasma/Cryptobia borreli* | | X | X | X |
| *Pharyngomonas kirbyi* | | | X | X |
| *Tsukubamonas globosa* | | | (X) | (X) |
| *Ploeotia vitrea* | | | (X) | (X) |
| All other species | | | | X |

Table Table S1: **Procedures that were applied to the data from species used in this study.** "X" means that the procedure was applied, "(X)" means that the procedure was applied, but species was excluded from the final proteome dataset, and blank space means the procedure was not applied.

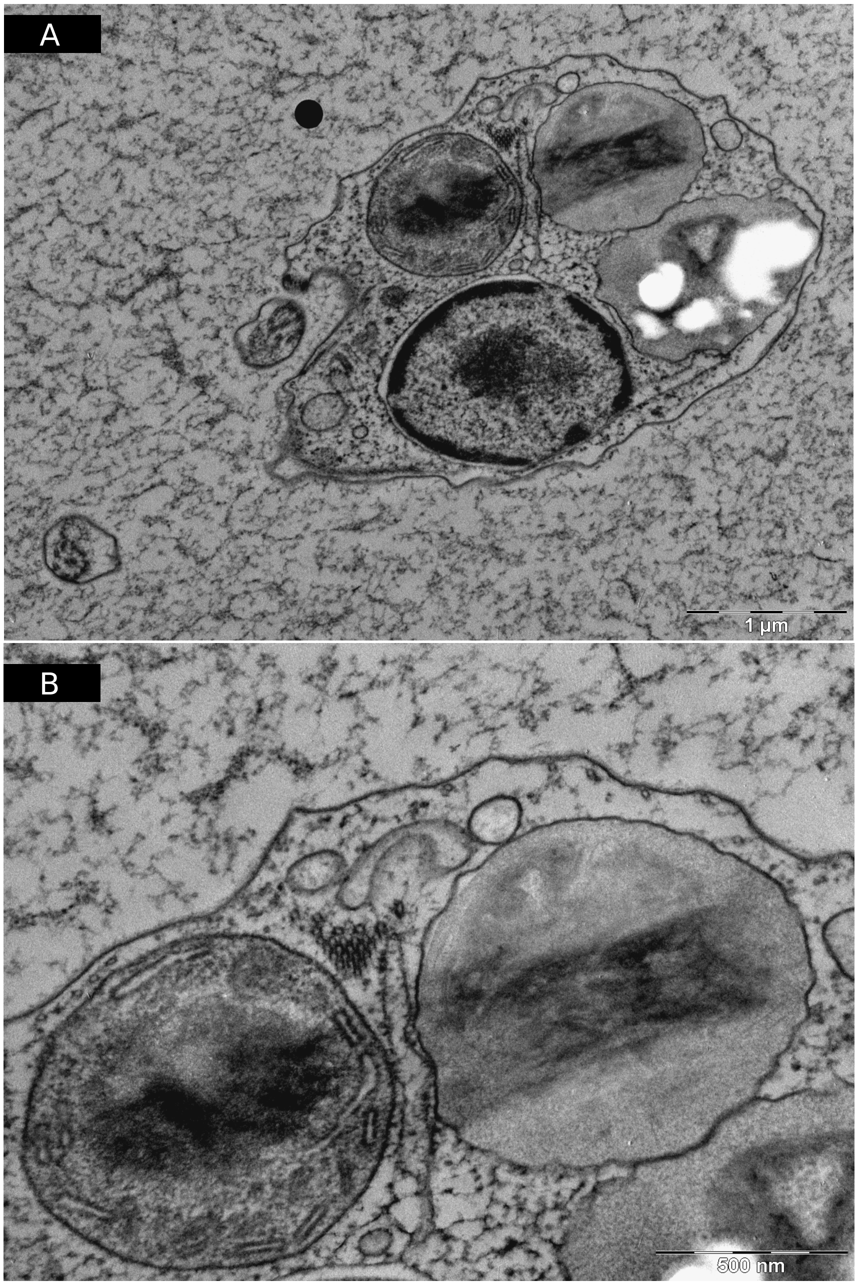


Figure Figure S2: **TEM image of 1D Neobodonid isolate BEAP0117-"13G".** Kinetoplast material wihin mitochondrion and "microtubular prism" are visible. All TEM Images of this isolate are available on FigShare `https://doi.org/10.6084/m9.figshare.30214630`.

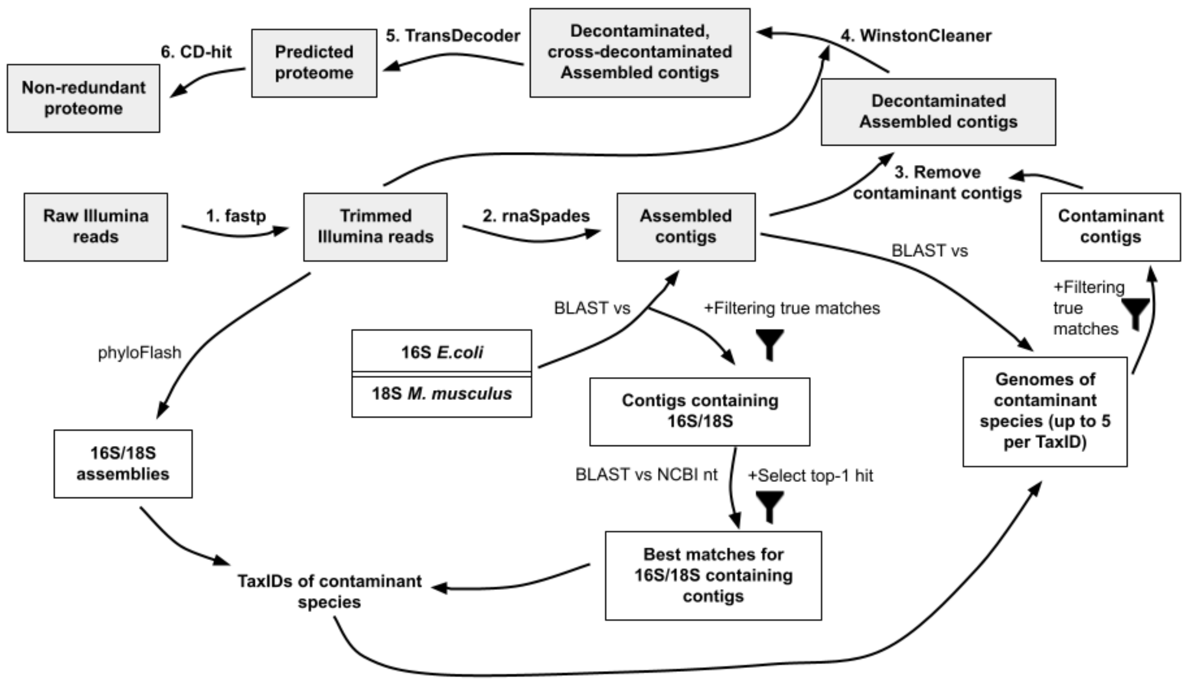


Figure Figure S3: **A graphical representation of transcriptome assembly, decontamination, protein prediction and redundancy reduction pipeline.** The numbers indicate the sequence in which different steps are performed. Shaded rectangles indicate the main data types produced throughout the process, while blank rectangles indicate data which was obtained for performing decontamination.

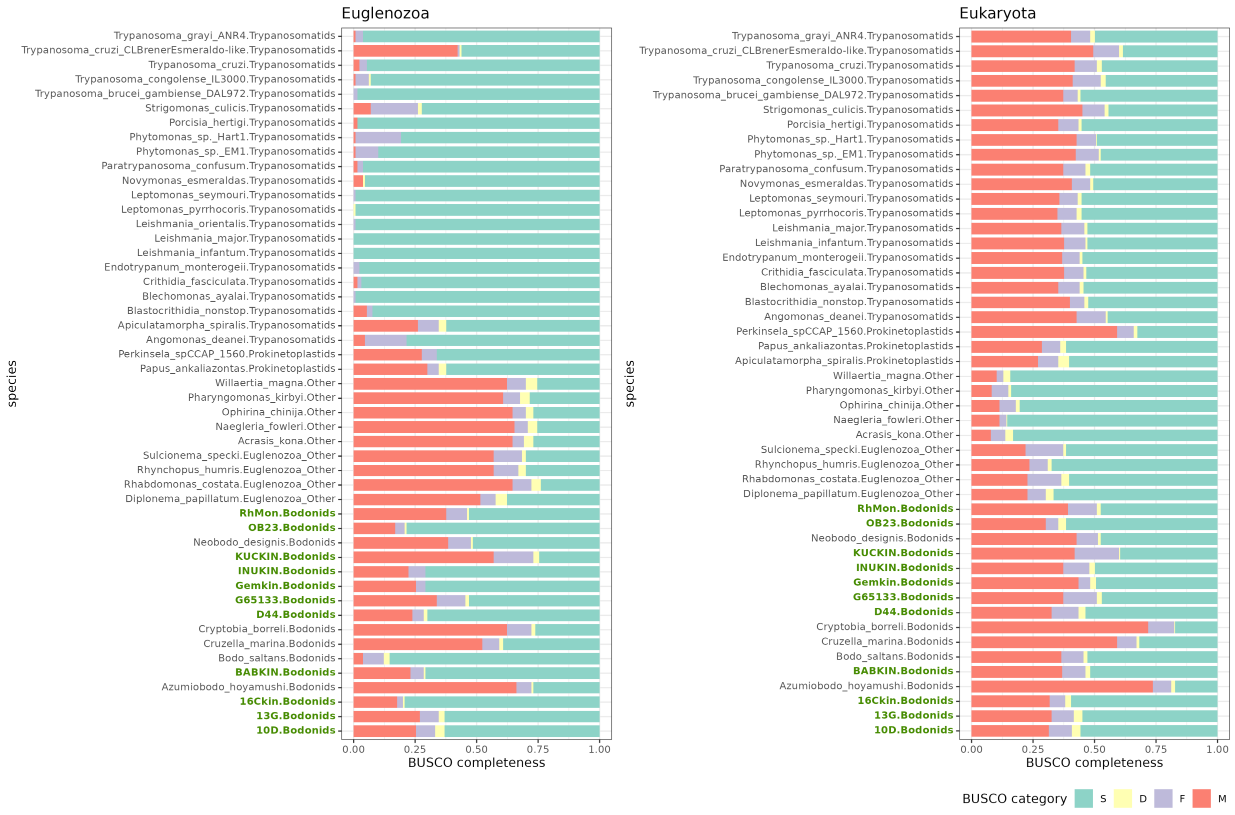


Figure Figure S4: **BUSCO scores of redundancy-reduced proteomes used in this study.** Completeness scores are shown for "Eukaryota" and "Euglenozoa" BUSCO v. 10 Manni et al. (2021) datasets. The x-axis represents the fraction of each BUSCO gene category. Colours of the bars indicate fractions of: "S" - Complete and single-copy BUSCOs; ("D" - Complete and duplicated BUSCOs; "F" Fragmented BUSCOs; "M" - Missing BUSCOs. On the y-axis, species and their taxonomic affiliation are indicated. isolates for which the data were produced in this study are highlighted in green.

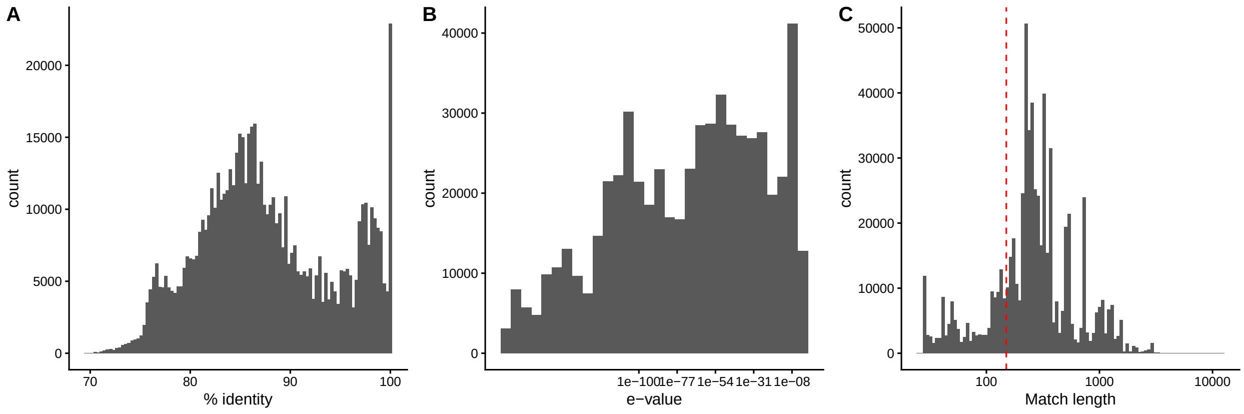


Figure Figure S5: **Distribution of BLAST hit statistics for the contigs of transcriptome assemblies generated in this study searched versus contaminant genomes.** 60,000 hits were randomly selected from all pairwise comparisons for visualisation purposes. The number of hits is represented on the vertical axis. On the horizontal axis: A - % identity of the hit, B - E-value of the hit, C - length of the match (the dashed vertical line represents the cutoff applied to hit match length). Minimum thresholds of % identity were selected individually for each pair of assembly and contaminant genome (see Methods); no threshold of E-value was applied. Note the log10 scales in B and C.

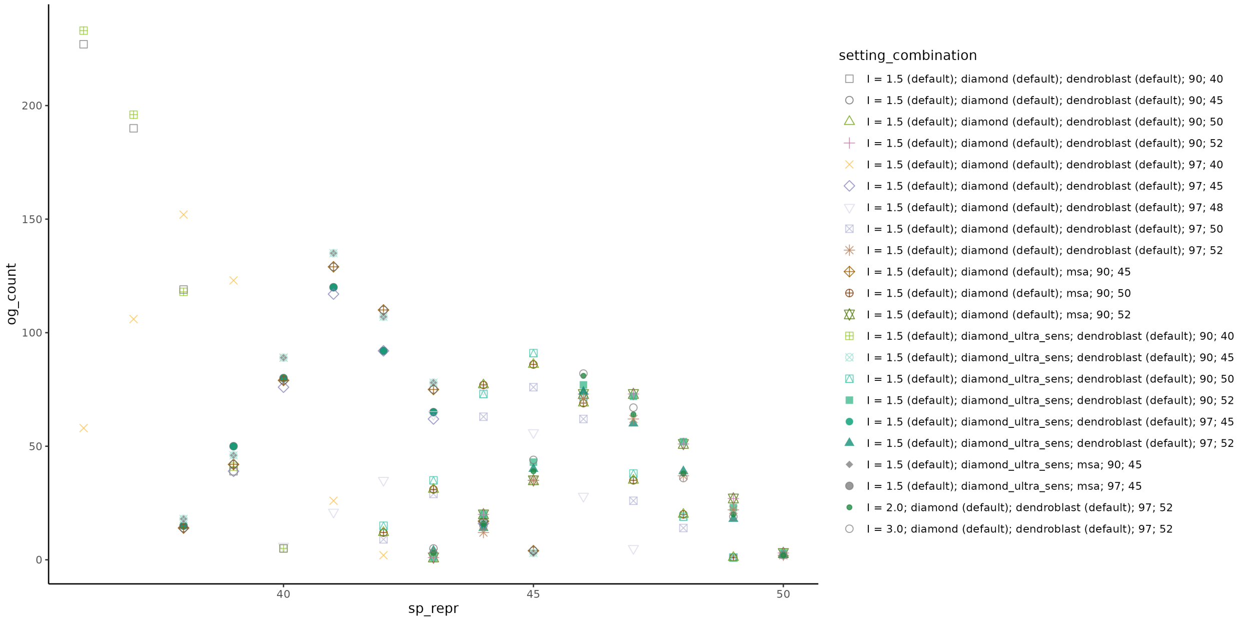


Figure Figure S6: **The number of OGs after subsetting for various setting combinations.** The vertical axis represents the count of OGs with the mean number of proteins per single species between or equal to 1 and 0.9 ("Mean", threshold value (a) as described in the text), standard deviation of number of proteins per single species lower than or equal to 0.5 ("SD", threshold value (b)), and the number of species with at least one protein in a given OG higher than 35 (">35 species", threshold value (c)). The horizontal axis represents the number of species with at least one gene in the OG. Shapes and colours of the points correspond to different combinations of settings in the following order: the OrthoFinder MCL inflation parameter; the OrthoFinder sequence search program; OrthoFinder method for gene tree inference; the CD-HIT amino acid global sequence identity cutoff; the number of taxa in the set.

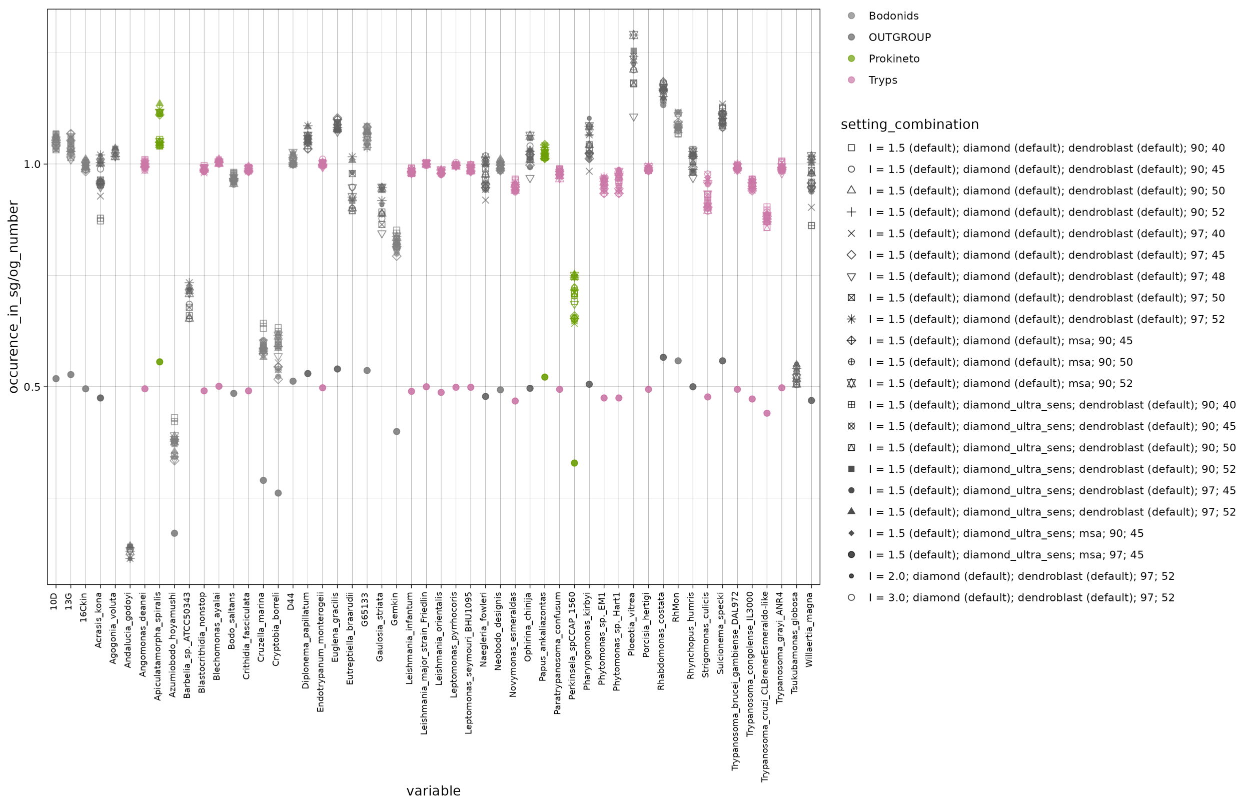


Figure Figure S7: **The occurrence of genes from different species in OG subsets for various setting combinations.** The vertical axis represents the fraction of OGs in the subset which contain at least one gene from a given species (listed on the horizontal axis). The shapes of points represent higher-rank taxonomic affiliation of a species: "Bodonids": Metakinetoplastina other than Trypanosomatida; "OUTGROUP": taxa selected as outgroup; "Prokineto": Prokinetoplastina; "Tryps": Trypanosomatida. Colours of the points correspond to different combinations of settings in the following order: the OrthoFinder MCL inflation parameter; the OrthoFinder sequence search program; OrthoFinder method for gene tree inference; the CD-HIT amino acid global sequence identity cutoff; the number of taxa in the set.

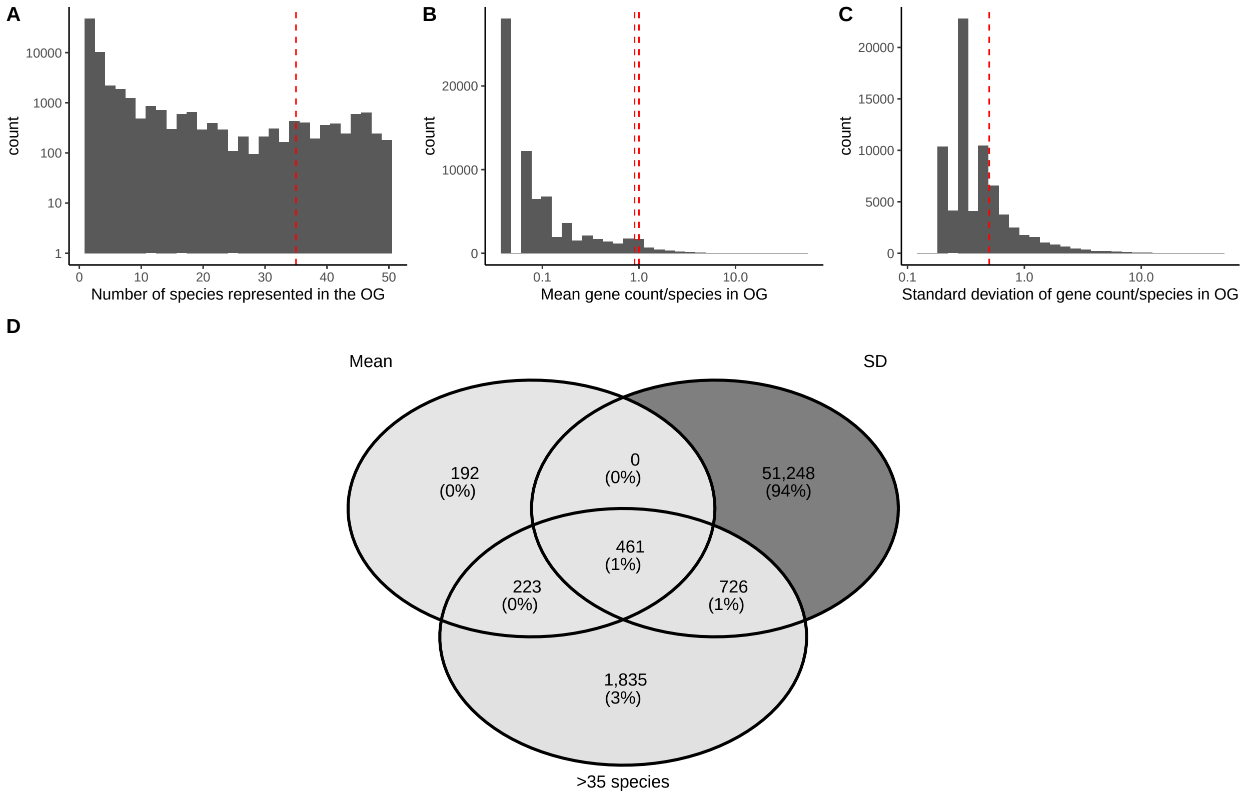


Figure Figure S8: **Subsetting OGs. A-C**: distributions of parameters used to establish thresholds to subset OGs. The count of OGs with a given parameter value is represented on the vertical axis, and the values of the three different parameters are represented on the horizontal axis. The dashed red vertical lines in A-C represent thresholds used for subsetting OGs. Note log10 scales in A-C. **D**: the count of OGs with the mean number of proteins per single species between or equal to 1 and 0.9 ("Mean"), standard deviation of number of proteins per single species lower than or equal to 0.5 ("SD"), and the number of species with at least one protein in a given OG higher than 35 (">35 species").

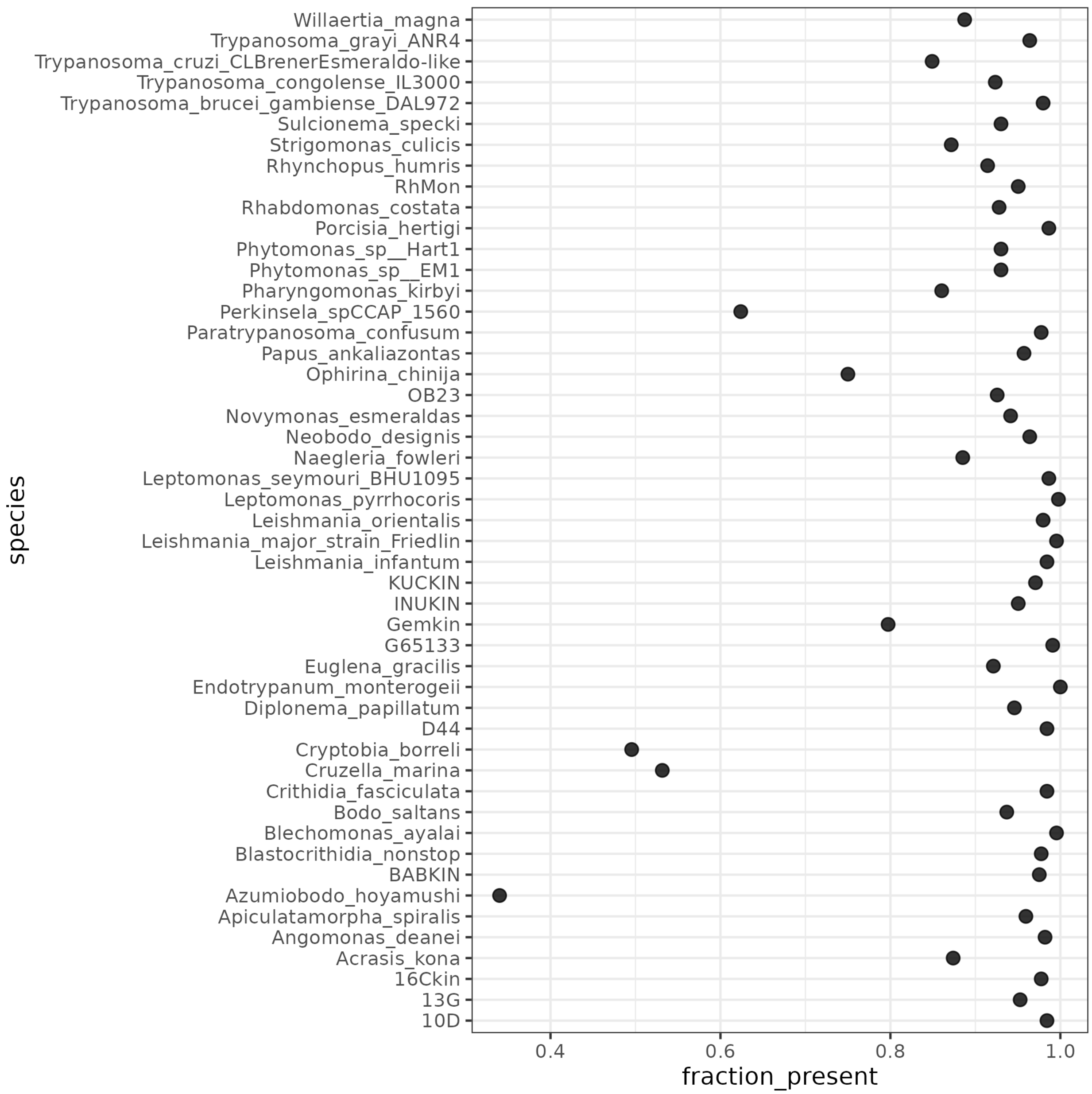


Figure Figure S9: **Fraction of OGs from the final OG dataset used in tree inference in which each species is represented.** The total OG count is 444.

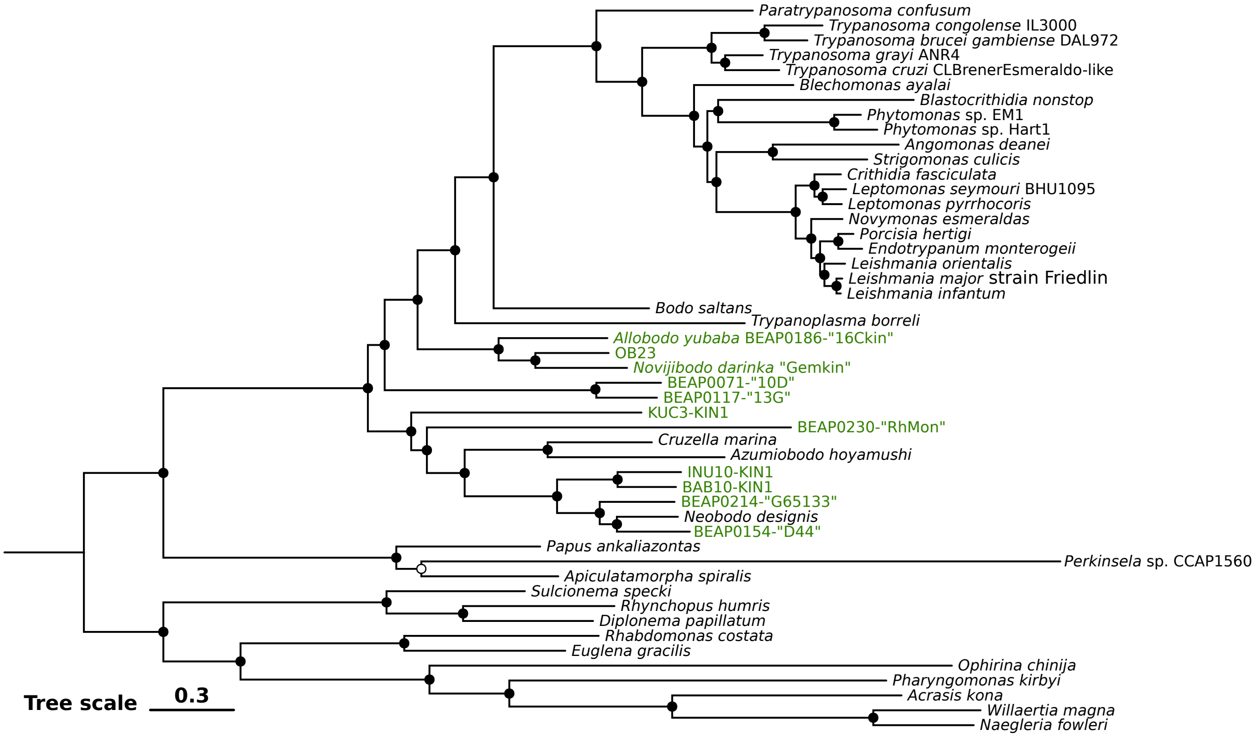


Figure Figure S10: **The topology of the Bayesian tree.** The tree represents the consensus topology of three independent chains run for approximately 16,000 generations each (with a burn-in of 4000 generations) with the CAT+GTR+G4 model, sampling once every generation. Posterior probability values at all nodes are 1, apart from the node labelled with a white dot, which has a posterior probability of 0.6. The scale represents substitutions per site. The tree topology differs from the topology presented in Figure 2 within the Prokinetoplastina, in which *Perkinsela* sp. is sister to *Papus ankaliazontas* + *Apiculatamorpha spiralis*. isolates for which the data were produced in this study are highlighted in green.
